# Colicin E1 opens its hinge to plug TolC

**DOI:** 10.1101/692251

**Authors:** S. Jimmy Budiardjo, Jacqueline J. Stevens, Anna L. Calkins, Ayotunde P. Ikujuni, Virangika K. Wimalasena, Emre Firlar, David A. Case, Julie S. Biteen, Jason T. Kaelber, Joanna S.G. Slusky

## Abstract

The double membrane architecture of Gram-negative bacteria forms a barrier that is effectively impermeable to extracellular threats. Bacteriocin proteins evolved to exploit the accessible, surface-exposed proteins embedded in the outer membrane to deliver cytotoxic cargo. Colicin E1 is a bacteriocin produced by, and lethal to, *Escherichia coli* that hijacks the outer membrane proteins TolC and BtuB to enter the cell. Here we capture the colicin E1 translocation domain inside its membrane receptor, TolC, by high-resolution cryoEM, the first reported structure of a bacteriocin bound to TolC. Colicin E1 binds stably to TolC as an open hinge through the TolC pore—an architectural rearrangement from colicin E1’s unbound conformation. This binding is stable in live *E. coli* cells as indicated by single-molecule fluorescence microscopy. Finally, colicin E1 fragments binding to TolC plugs the channel, inhibiting its native efflux function as an antibiotic efflux pump and heightening susceptibility to three antibiotic classes. In addition to demonstrating that these protein fragments are useful starting points for developing novel antibiotic potentiators, this method could be expanded to other colicins to inhibit other outer membrane protein functions.

## Introduction

In Gram-negative bacteria, the concentric structures of the outer membrane, cell wall, and cytoplasmic membrane protect the cell from extracellular threats. Of these protective structures, the outer membrane forms a particularly formidable barrier(*5*), owing to the impermeability of the lipopolysaccharide (LPS) layer that constitutes the outer membrane(*7*). The primary means by which external molecules can gain access to the cell is through the ∼100 varieties of barrel-shaped proteins that are embedded in each bacterium outer membrane(*8*) and whose diverse functions include the transport of molecules across the membrane—specifically, the import of nutrients and metabolites and the export of toxins and waste.

Because outer membrane proteins (OMPs) are accessible from outside the cell, bacteriophage and bacterial toxins have evolved to exploit OMPs to initiate delivering cargo across the outer membrane. Bacteriocins hijack the OMPs of a target bacterium to cross its impermeable outer membrane and kill the bacterium. Colicins are *E. coli*-specific bacteriocins, protein toxin systems through which bacteria engage in bacterial warfare with other, similar bacteria. Although colicins differ in their receptor targets and killing mechanisms, most colicins share a common tri-domain architecture, comprising the following components: (i) an N-terminal translocation (T) domain, (ii) a receptor-binding (R) domain, and (iii) a C-terminal cytotoxic (C) domain (Figure 1A). Much of what is known of E colicin import has been determined through studies of the colicin E3 and E9 as reviewed by Cramer et al.(*9*) Import is initialized by R domain binding to the vitamin B12 transporter, BtuB, with high affinity(*10, 11*); this binding localizes the colicin onto the outer membrane. Once the colicin is tethered to the outer membrane surface, the T domain initiates translocation using the secondary OMP receptor OmpF to access TolA/Pal system for group A colicins or the Ton system group B colicins. In most cases, the T domain requires an OMP distinct from the R domain target.(*12*) A handful of colicins have been structurally characterized with their OMP counterparts, although not in their entirety. Previous structures of short N-terminal fragments of the T domain of ColE9 with OmpF(*13*) and the The R domains of ColE2/E3 and Ia with BtuB(*11, 14*) and Cir(*10*), respectively showed that T domains fully penetrate deeply into lumen of the outer membrane receptors while R domains interact with the extracellular loop regions of their receptors. Studies of the bacteriocin pyocin S5 from *Pseudomonas aeruginosa* suggest that bacteriocin architectures and mechanisms may be conserved across all Gram-negative species.

**Figure 1.**
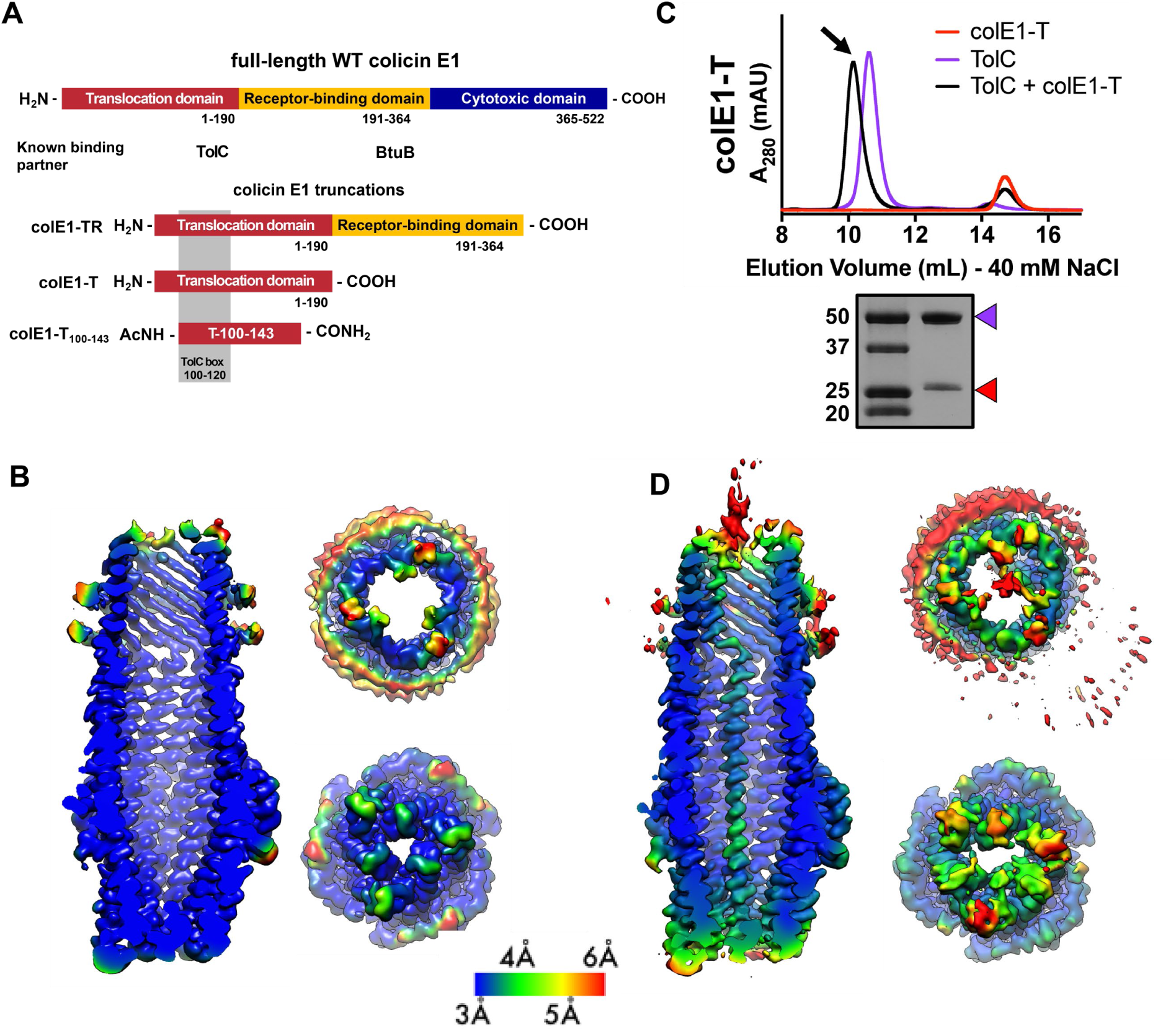
Cell-surface localization of colicin E1 fragments. (**A**) Architecture of full-length colicin E1 showing domains and their known binding partners. Three truncation constructs used in this study (**B**) CryoEM structure of TolC embedded in nanodiscs. Side, top, and bottom views are colored by local resolution, as computed in cryoSPARC from the final half-maps. The side view is cropped to display the particle interior. (**C**) SEC chromatogram of colE1-T (red line) and TolC (purple line). The arrow indicates the co-elution (black line) fractions that were analyzed by SDS-PAGE. On the SDS-PAGE gel (bottom), red arrows indicate the presence of the colicin E1 construct that has co-eluted with TolC. (**D**) The CryoEM structure of colE1-T bound to TolC and colored by local resolution as in (B).

Colicin E1 is a bacteriocin secreted by *E. coli* that enters the periplasm of neighboring cells and forms a pore on the cytoplasmic membrane leading to membrane depolarization and cell death. Colicin E1 uses TolC, the outer membrane component of the acridine efflux pump, as the T domain receptor(*15*) and BtuB as the R domain receptor(*16*). A high-resolution x-ray structure exists of only the cytotoxic domain of colicin E1(*17*) but not the T or R domains, thus no mechanistic insights can be gained about initial steps of import. Domain swapping experiments (*15*) between colicin A and E1 identified regions of the protein that interact with specific outer membrane proteins. Early studies of pore forming colicins, including E1, showed that cytotoxic-domain-induced K^+^ efflux on the inner membrane can be reversed after it has begun by subsequent addition of trypsin to cells. This, in addition to trypsin’s inability to cross the outer membrane, led to the belief that portions of colicin remain tethered to their OMP partners as the cytotoxic domain depolarizes the cytoplasmic membrane.(*18–21*)

In order to understand the early stages of colicin E1 import, functional studies of truncations of colicin E1 which lack the cytotoxic domain have been characterized *in vitro*. Residues 100-120 of colicin E1 (termed the ‘TolC box’, Figure 1A) have been shown to be necessary but not sufficient for binding TolC. Peptides that include the TolC box co-elute with TolC(*22*) and disrupt channel conductance(*22, 23*). Moreover, *E. coli* exposure to TolC-box-containing peptides can prevent subsequent binding and cytotoxicity of full-length colicin E1(*24*). Circular dichroism of the T domain indicates that it exists as a helical hairpin (closed hinge) in solution similar to other colicin T domains(*25*) and that the proline at the center of the TolC box forms its apex(*22*). This measurement led to the proposal, known as the “pillar model,”(*22*) that the T domain inserts into the TolC barrel as a helical hairpin where the N and C-termini are pointing to the cell exterior. According to this model, the hairpin stuck in TolC acts as a buttress to facilitate the cytotoxic domain entry directly through the membrane.

A competing model for colicin E1 import, known as the “total thread” model(*9, 22*), posits that the protein unfolds and passes through TolC N-terminus-first as an unstructured peptide, followed by refolding in the periplasm. In this model the binding between the intrinsically unstructured colicin N-termini and periplasmic proteins(*24, 26*) creates a pulling force that results in the translocation of the whole colicin.

Here we use cryogenic electron microscopy (cryoEM) to solve the first high-resolution structure of a bacteriocin, colicin E1, bound to TolC (*27*). We find that colicin E1 binds stably to TolC, not as a helical hairpin but as a single-pass folded helix with the N-terminus inside the periplasm. Additionally, we find that the colE1 TR domain binds TolC *in vivo* as well. Using single-molecule fluorescence microscopy, we find that colE1-TR, lacking the cytotoxic domain, remains stalled on the outer membrane and does not fully translocate into cells. Lastly, we leverage this stalling of ColE1 to block the native TolC function as an antibiotic efflux pump. Because they are accessible from outside the cell, OMPs are attractive targets for the development of novel antibiotics, and research has begun to reveal the therapeutic potential of interfering with OMP structure and function. Recently, a monoclonal antibody was found to inhibit OMP folding with bactericidal effects(*28*).

Here by determining the structure and mechanism of colE1 insertion we establish an alternative approach for targeting OMPs—the development of molecular plugs that block OMP pores. Such plugs would allow for the manipulation of bacterial transport, providing a means of either starving the bacterium by preventing the import of valuable nutrients or poisoning the bacterium by preventing the export of toxins. Through real-time efflux assays, minimum inhibitory concentration (MIC) experiments, we find that colE1-TR and colE1-T are able to inhibit TolC-mediated efflux. We find that this plugging of TolC reduces the amount of antibiotics required to inhibit the bacterial growth—indicating that this colicin E1 fragment reduces the antibiotic resistance conferred by TolC.

## Results

### CryoEM structure of TolC embedded in nanodiscs

To determine the structural details of colicin E1 binding to TolC, we solved the cryoEM (cryogenic electron microscopy) structure of TolC embedded in nanodiscs with and without added colE1-T (PDB 6WXH and 6WXI, respectively). We recently reported a high-yield TolC preparation by refolding from inclusion bodies(*29*) and inserted refolded TolC into nanodiscs. The cryoEM structure of refolded TolC alone (Figure 1B) is similar to the previously published crystal structures of natively derived TolC alone(*30*) or in complex with hexamminecobalt(*31*) but with more splayed loops at the periplasmic opening (residues 165-175). There was local variation in resolution within the model with lower resolution associated with the extracellular/periplasmic ends and the nanodisc scaffold protein (Figure 1B). The lower residue resolution at these apertures may indicate dynamics not captured in the x-ray crystal structures.

We next determined binding of colicin E1-T domain to capture the complex for structure determination. Residues 1-190 (colE1-T) which span the translocation domain and are known to bind to TolC (Figure 1A). We determined colE1-T:TolC binding *in vitro* via co-elution by size exclusion chromatography (SEC) as previously described(*6*). When colE1-T and TolC were mixed, we observed a leftward shift in the TolC peak and a decrease in intensity associated with the colE1-T peak indicating that a subset of the population has migrated with TolC (Figure 1C). We analyzed the peak (Figure 1C arrow) using SDS-PAGE and found the presence of both colE1-T and TolC. The unimodal shifted peak observed with colE1-T:TolC indicates that there is a single species of fully bound TolC. When resolved by cryoEM, addition of colE1-T to TolC breaks the three-fold channel symmetry and the additional protein is observed running all the way through the TolC barrel as a single-pass, all-α-helical chain spanning more than 130 Å (Figure 1D). The maps refined to nominal global resolutions of 2.84 Å and 3.09 Å for the TolC and colE1-T:TolC, respectively (Table S1).

The colE1 T domain inserts into TolC with its amino terminus pointing inwards through the periplasmic iris, and 2D class-averages show that the carboxy terminus of the helix continues and projects out into the extracellular space (Figure 2A). 85 of the 190 colE1-T residues could be modeled to this density (residues 46-131) (Figure 2B). No such regular protrusion was seen for the glycine-rich colE1 amino-terminus, which we expect to be disordered in the periplasmic space(*22, 26, 32*). The colE1 chain binds TolC asymmetrically, primarily contacting only one of the three TolC chains.

**Figure 2.**
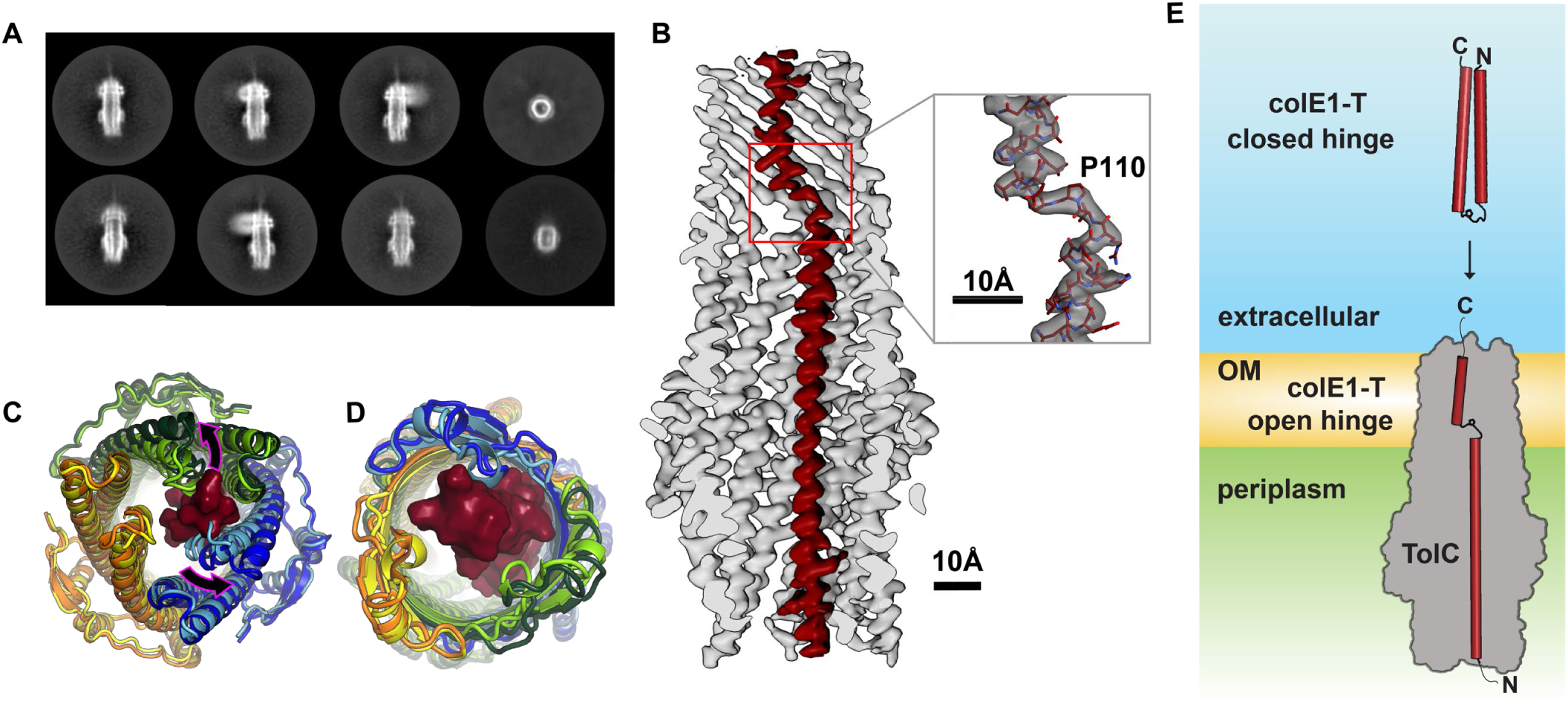
Colicin E1 binds to TolC as a single-pass kinked helix. (**A**) 2D class averages of colE1-T bound to TolC imbedded in nanodiscs (**B**) Cutaway view of the TolC interior (gray). ColE1-T (red) binds asymmetrically in an open-hinge conformation. The hinge region of colE1T including P110 shown in stick representation (inset). (**C** and **D**) The dilated terminal apertures. The apo cryoEM structure of TolC (subunits colored light blue, light green, and yellow) compared to holo cryoEM structure (subunits colored dark blue, dark green and orange). (**C**) Periplasmic aperture. (**D**) Extracellular aperture. (**E**) In the unbound state, colE1-T (red cylinders) exists as a closed hinge. When bound, colE1-T is in an open-hinge conformation through TolC with the N-terminus into the pore of TolC (grey). The homology model of colE1-T in its solution state was built using I-TASSER(*4*) and is consistent with previous experiments(*6*).

Compared to the unbound state, asymmetric colE1 binding dilates the periplasmic TolC aperture (Figure 2C) and, to a lesser extent, the extracellular aperture (Figure 2D) so that they can accommodate colE1 in the absence of any other proteins or motive force. Although the terminal apertures dilate and are filled by the ligand and the TolC box forms specific interactions with the TolC inner wall, the large internal volume of TolC is not fully occupied by colE1.

In solution, unbound colicin E1 is a two-helix hairpin as indicated by far UV circular dichroism (CD)(*6*). A homology model built using I-TASSER(*4*) also predicts the closed hinge conformation with proline 110 at the apex (Figure 2E top). However, our cryoEM reconstruction of colicin E1 in complex with TolC shows that colE1-T hinges open at the TolC box to an extended conformation upon binding to TolC (Figure 2B, and 2E bottom). The colE1-T helices have a short kink in the middle around proline 110, precisely at the proposed turn location in the hairpin model (Figure 2B inset). This area is also the center of the TolC box that has been known to be critical for the colE1:TolC interaction(*22, 24*).

### Colicin E1 makes residue specific interactions with TolC

TolC forms a large rigid conduit in the outer membrane and periplasm (Figure 3A) with a water filled lumen. The hydrophilic nature of the channel(*33*) coupled with the inflexibility of the outer-membrane-embedded barrel largely precludes the formation of any hydrophobic interfaces that are typically the basis of protein-protein interfaces involving helical peptides. Yet, we did observe specific polar contacts between colicin E1’s TolC box and the TolC barrel.

**Figure 3.**
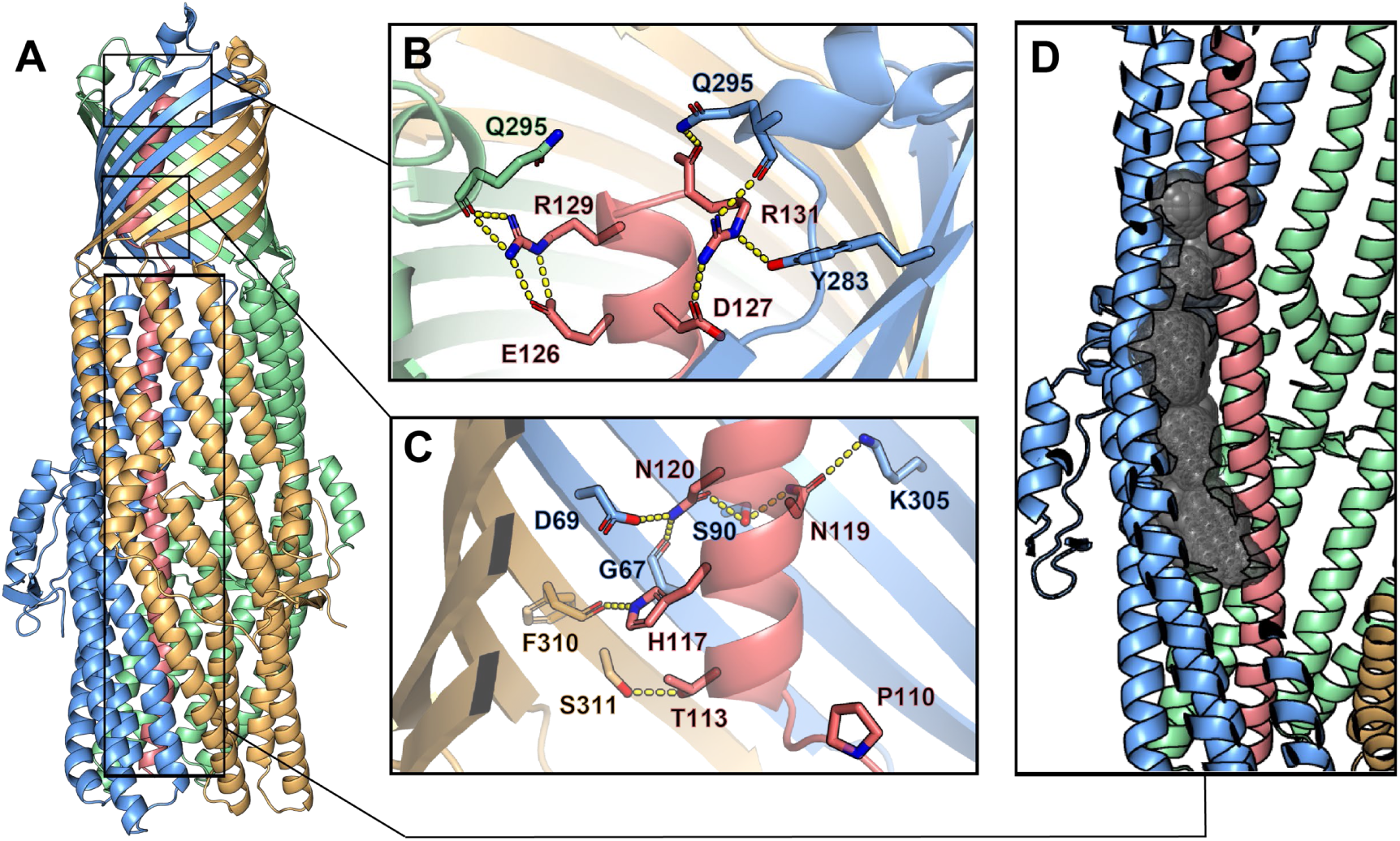
Inter-chain contacts between colE1T and TolC. (**A**) Molecular dynamics simulation refined structure of the colE1-T:TolC complex colored by chain (colE1-T: light red, TolC: light blue, light green, light gold) (**B-C**) Polar interaction network of colE1-T and TolC on the C-terminal side of the proline kink in the β-barrel region. (**B**) near the extracellular opening of TolC and (**C**) before the transition to the TolC periplasmic helical region. Residues involved in forming polar interactions and proline 110 are shown in sticks. (**D**) ColE1-T spanning the TolC helical barrel does not make full contact with the side of the TolC barrel. The cavity (black spheres) between colicin (light red) and one chain of TolC (light blue) was detected using GHECOM(*1–3*)

To obtain the most accurate atomic model for interpretation of atomic interactions between peptides, we utilized map-restrained molecular dynamics in model refinement(*34*). The refined model had improved chemical plausibility (for instance, MolProbity(*35*) score improved from 1.25 to 0.50) and polar contact were more easily identified. Because this refinement improved the concordance between the map and model, we conclude that the use of cryoEM density as a restraint was successful in preventing overfitting. Specifically, the EMRinger(*36*) score improved from 2.77 to 3.81 and the map-model FSC=0.5 improved from 3.44Å to 3.37Å.

The atomic model showed significant polar interactions form between colicin E1 and TolC at the apertures and around the TolC box. The acidic patch of colicin E1 near the extracellular aperture contains two arginine residues (R129 and R131) that form hydrogen bond networks with one TolC chain each (Figure 3B). In addition, colicin E1 residues T113, A116, N119, N120 (all part of the TolC box) establish polar interactions with TolC residues G67, D69, S90 and K305 on one TolC chain (Figure 3C, light red and light blue) while colicin H117 interacts with TolC F310 on a neighboring TolC chain (Figure 3C, light red and light gold). By contrast with the β-barrel of TolC, in the interior of periplasmic helical barrel of TolC, colicin E1 does not make full contact with the α-helical barrel interior and there is a gap between the two proteins (Figure 3D). This is in agreement with previous studies that showed a colicin truncation (residues 1-100) that ends just before the beginning of the TolC box does not bind to TolC(*22*) or prevent cytotoxicity of full-length colicin in cells(*24*).

Moreover, this structure is consistent with reports that mutations at TolC sites G43 or S257 abrogate colE1 activity(*37*) as these residues are contact sites between TolC and colE1 (Figure S1). Conversely, the absence of any effect of mutating colicin R103 and R108(*24*), is consistent with interaction those residues have with the barrel solvent rather than TolC. The newly solved structure, in combination with previous work(*22, 24*), indicates the specificity of colicin E1 binding to TolC is encoded in the portion that binds to TolC β-barrel.

### Membrane localization of colE1 truncations

Since we find the colE1 T domain binds to TolC, we investigated if colicin E1 fragments bind to the native TolC in *E. coli* cells. Though ColE1 C domain activity—depolarizing the inner membrane—requires the C domain to pass through the outer membrane(*38*), it remains unclear if the colE1-TR domains translocate as well. We first determined *in vitro* binding of colE1-TR, which includes residues 1-364 (colE1-TR) (Figure 1A) which contains the N-terminal portion that binds to TolC and the C-terminal portion that binds to BtuB. ColE1-TR aggregates at the salt concentration (40 mM) used for the co-elution experiment used for colE1-T so we increased the NaCl concentration to 200 mM. ColE1-T still binds to TolC at 200 mM NaCl although to a much lesser degree (Figure S2). Unlike colE1-T (Figure 1C), colE1-TR shows a bimodal distribution indicative of a mixed population of bound and unbound TolC (Figure 4A). Using an extracellular protease digestion assay, we assessed whether colE1-T and colE1-TR translocate into the cell or localize on the cell surface. ColE1 fragments were incubated with cells, washed, and treated with or without trypsin to digest any extracellularly bound protein to cells. When probing for the C-terminal epitope by immunoblotting, there was no detectible colE1-T binding (Figure 4B, left), though there was detectible binding of ColE1-TR (Figure 4B, right). Moreover, we found that if colE1-TR is incubated with cells and subsequently treated with increasing trypsin concentrations, the colE1-TR band disappeared, indicating that the colicin E1 fragment was localized to the outer membrane surface (Figure 4B, right). The periplasmic control SurA was not degraded at any trypsin concentration unless the cells were lysed before trypsin digestion(*39*).

**Figure 4.**
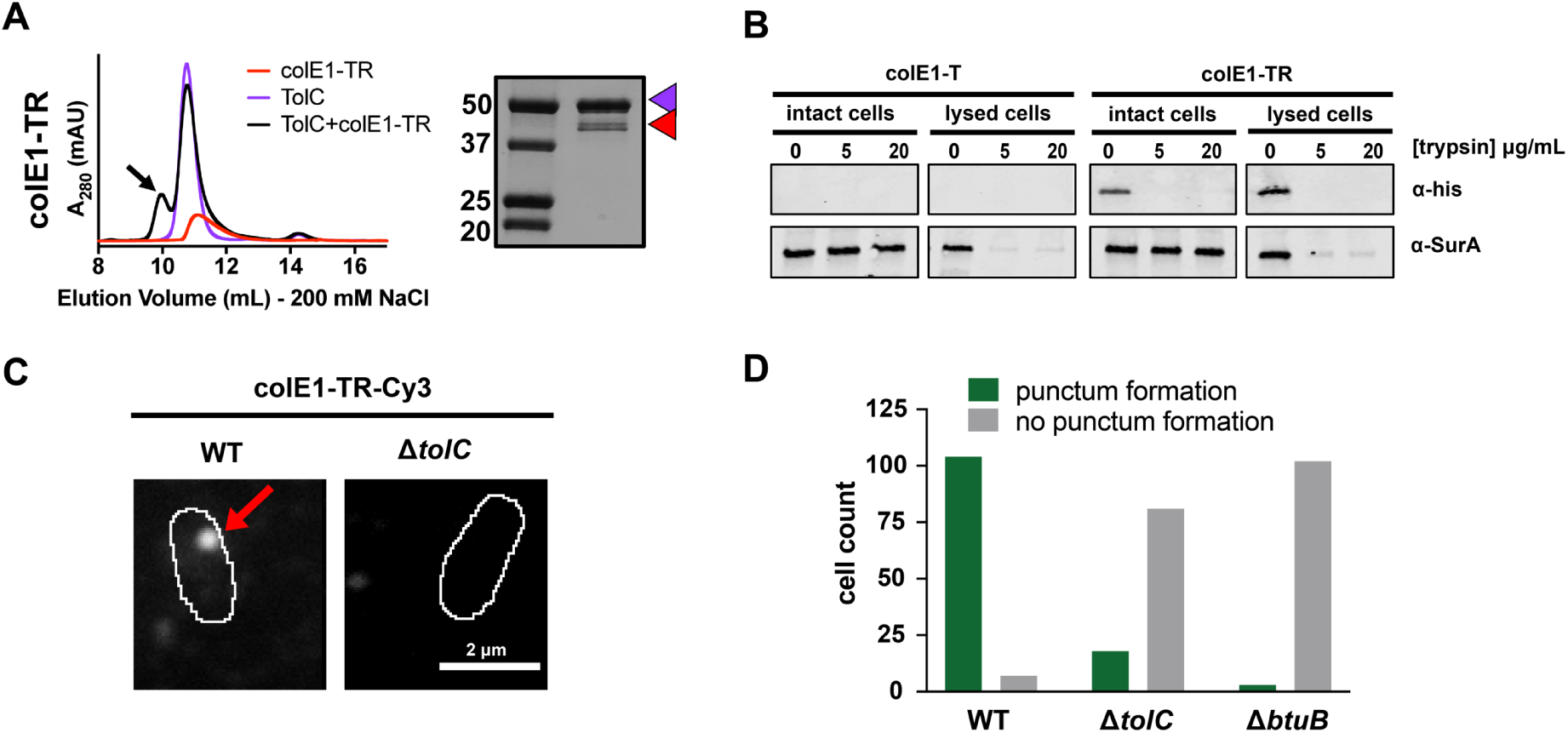
Colicin E1-TR localizes on the outside of the cell. (**A**) SEC chromatogram of colE1-TR (red line) and TolC (purple line). The arrow indicates the co-elution (black line) fractions that were analyzed by SDS-PAGE. On the SDS-PAGE gels (right), red arrow indicates the presence of colicin E1 construct that has co-eluted with TolC (purple arrow). (**B**) Extracellular protease digestion assay with two colicin E1 truncation constructs, each labeled with a C-terminal His-Tag. Periplasmic SurA was used as a membrane integrity control. (**C**) Fluorescence image of colE1-TR-Cy3 overlaid on outlines of living *E. coli* cells from phase-contrast microscopy for WT and Δ*tolC*. Red arrow points to a punctum. (**D**) Cell counts where colE1-TR-Cy3 punctum formation was observed for WT, Δ*tolC*, and Δ*btuB*. Number of cells observed, *n* = 111, 91, 105, respectively.

Since the proteolysis experiments are not sensitive enough to detect if only a few molecules have entered the cell, we further probed the interaction between the cell membrane and surface-localized colE1-TR with single-molecule fluorescence microscopy using the fluorescent dye Cy3. This method is able to detect single molecules on the cell surface as well as those within the cell indicative of those molecules having crossed the outer membrane(*40, 41*). When colE1-TR-Cy3 was added to the extracellular environment of WT BW25113 *E. coli* (containing TolC), distinct puncta (Figure 4C, left) formed on 94% of the cells (number of cells, *n* = 111) (Figure 4D); cells with observed puncta most often featured a single punctum, though a small fraction had 2 puncta. On average, the WT cells had 1.2 puncta. In a Δ*tolC* strain, puncta were observed on only 18% of cells (*n* = 99) (Figure 4C, right; Figure 4D); on average, the Δ*tolC* cells had 0.2 puncta. As a Δ*btuB* control strain, we used BL21 (DE3) which is known to have a premature stop codon at *btuB* residue 58(*42*). Puncta were observed on only 3% of cells lacking BtuB (*n* = 105) (Figure 4D and S3A); on average, the Δ*btuB* cells had 0 puncta. In WT and Δ*tolC* cells that featured puncta, colE1-TR formed puncta consistent with ∼20 molecules based on dividing the puncta intensity by the fluorescence intensity of the last fluorescent molecule before photobleaching (*43*). The observed size and number of molecules are in agreement with previous studies of BtuB clusters (*44, 45*) and the punctum locations within the cell were variable. To rule out punctum formation as an artifact of Cy3 conjugation, we found colE1-TR-GFP displayed the same cluster formation as colE1-TR-Cy3 (Figure S3B). No other single protein binding events were detected aside from the puncta on either WT or Δ*tolC* cells.

Fluorescently labeled pyocins, the colicin analog in *Pseudomonas*, have previously been used to detect translocation across the outer membrane of *Pseudomonas aeruginosa*(*46*). Here we use an analogous experiment with fluorescently labeled colE1 to determine cellular localization in *E. coli*. In time courses, bound colE1-TR puncta remained immobile for > 5 minutes (Movie S1 corresponds to the first 8 seconds of data used to attain the WT image in Figure 4C). This is consistent with continued association of colE1-TR with membrane-embedded BtuB, which has limited mobility(*47*). This result indicates that colE1-TR does not fully translocate(*46*), because if colE1-TR entered the periplasmic space it would freely diffuse on these timescales.

Colicin constructs lacking the R domain (colE1-T-Cy3) showed no detectable binding either to WT or Δ*tolC* cells (Figure S3C), indicating that the TolC-colE1-T interaction is much weaker than the BtuB-colE1-TR interaction.

Because we only see one punctum per cell, we anticipate that some BtuB and TolC remain unbound because of the geometric constraints of punctum formation. Though BtuB and TolC need to be in close proximity for colE1 binding to occur, when BtuB clusters together in groups of 12 or more, it may exclude TolC. These cluster geometries would therefore lower the number of full binding sites available for the T and R domains of colicin E1.

### ColE1-TR inhibits efflux and makes *E. coli* more susceptible to antibiotics

Since the colicin truncations are stalled on their respective OMP receptors, we investigated if this stalling could disrupt native TolC efflux. Real-time efflux inhibition by colicin E1 fragments was assessed using a live-cell assay with N-(2-naphthyl)-1-naphthylamine (NNN)- dye, which is effluxed by the acridine efflux pump and fluoresces when it is localized inside the cell(*48*). NNN efflux can be turned off by the protonophore CCCP, which neutralizes the proton motive force and allows NNN to accumulate within the cell. Active efflux can then be monitored by the fluorescence decay once proton motive force is reenergized by glucose addition(*48–52*).

We assessed the ability of colicin E1 truncations to plug TolC by monitoring real-time NNN efflux. Cells exposed to colE1-T_100-143_ did not show lower real-time efflux than untreated cells (Figure 5A). This observation is notable in light of the fact that, in previous conductance studies, similar peptides were shown to bind TolC(*6, 24*) and to occlude the channel(*6, 23*).

**Figure 5.**
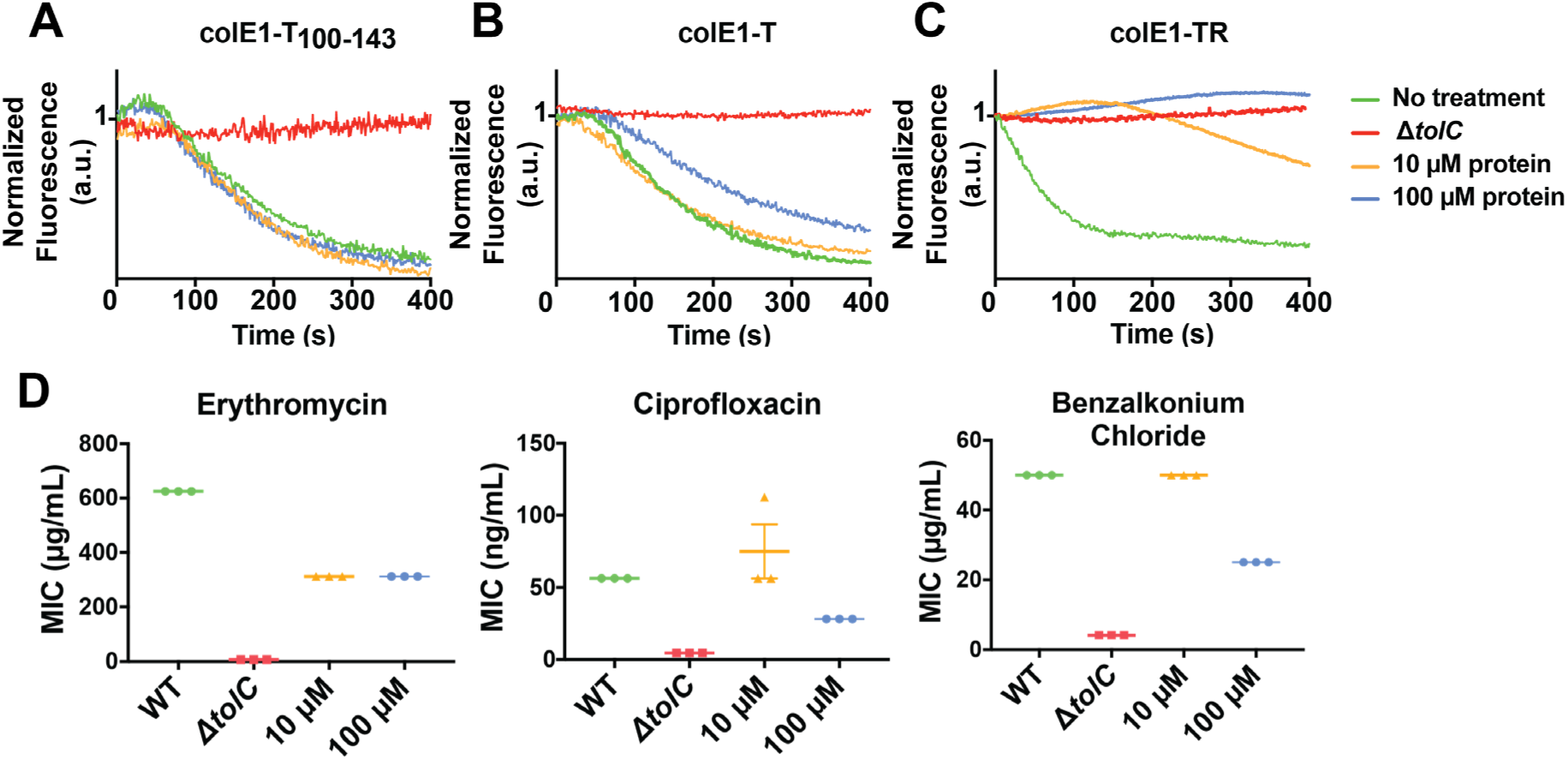
Colicin E1 fragments inhibit efflux and potentiate antibiotics. (**A-C**) Effect of colicin E1 fragments on efflux: WT with no protein (green), Δ*tolC* (red), WT + 10 μM (orange), WT + 100 μM protein (blue). (**A**) colE1-T_100-143_, (**B**) colE1-T, (**C**) colE1-TR. (**D**) Antibiotic susceptibilities in the absence (green) and presence of colE1-TR added to WT at 10 μM (orange) and 100 μM (blue). MICs for Δ*tolC* (red) are included as a reference. Data are biological replicates reported as the mean and individual data points.

Cells exposed to colE1-T showed less decay in final fluorescence than untreated cells, indicating that colE1-T partially inhibits efflux (Figure 5B). Finally, exposure to colE1-TR produced no decrease in fluorescence, showing full inhibition of efflux (Figure 5C).

Because colE1-TR completely inhibits NNN efflux, we evaluated the capacity of colE1-TR to potentiate antibiotics representing three different classes that are known TolC substrates: ciprofloxacin, erythromycin, and benzalkonium chloride (from fluoroquinolones, macrolides, and quaternary ammonium compounds, respectively). An effective TolC plug will reduce the concentration required to inhibit growth as antibiotics remain trapped within the cell. WT *E. coli* cells that were exposed to 100 μM colE1-TR with each of these antibiotics showed significantly lower minimum inhibitory concentrations (MICs), the lowest concentration of antibiotic that inhibits visible growth, than cells exposed to the antibiotics alone (Figure 5D): exposure to 100 μM colE1-TR made WT *E. coli* ∼2 to 7-fold more susceptible to these antibiotics (Table S2). Exposure of WT cells to 10μM colE1-TR was sufficient to potentiate erythromycin. Engagement of TolA by the N-terminal TolA box of colicin E9 has previously been shown to cause outer membrane defects(*53*) which would lead to enhanced antibiotic susceptibility. To rule out outer membrane-mediated defects caused by the TolA box in colE1, we created a truncation of colE1-TR (colE1-TR_Δ1-40_) lacking the N-terminal 40 residues including the TolA box. The MIC of colE1-TR and colE1-TR_Δ1-40_ are identical (Figure S4) indicating that colE1-TR engagement of TolA does not contribute to the observed antibiotic susceptibility.

## Discussion

There are two conflicting models of colE1 translocation: 1) the total thread model in which the entire colicin unfolds and threads through TolC(*24, 54*) and 2) the pillar model, in which colicin E1 inserts into TolC as a helical hairpin, facilitating LPS-mediated self-translocation of the colicin cytotoxic domain(*6*). Our data support aspects of both models. The belief in a closed hinge conformation of bound colE1 prompted two arguments against the total thread mechanism: 1) the colE1-T closed hinge conformation is too wide to fit through TolC and 2) both ends would face the extracellular milieu(*6, 54*). The unanticipated colE1-T hinge opening in the bound state resolves these objections: colicin threads into the TolC barrel with the amino terminus pointing into the periplasm, as the total thread model hypothesized. However, our trypsin digests and single-molecule fluorescence images show colE1-TR stalls at the outer membrane. Though we cannot exclude the possibility that full length protein translocation depends on the C domain, we observe—as hypothesized by the pillar model—that colicin sits inside TolC but does not translocate. However, the architecture of the complex differs from that hypothesized by the pillar model. These results are in agreement with early studies of pore-forming colicins in which trypsin added to the extracellular environment can reverse colicin activity. Digestibility by trypsin, as well as the ability of colE1-T/TolC to reconstitute a monodisperse stable complex *in vitro* are consistent with the colicin remain in in contact with its outer membrane receptors while the cytotoxic domain depolarizes the inner membrane.

In addition to demonstrating a colicin insertion mechanism, our observations form the basis of a means to manipulate bacterial import and efflux. Though colicin E1 confers some level of antibiotic potentiation, the relatively high concentration of colE1-TR needed to inhibit efflux may be explained by a combination of the geometric constraints of creating large clusters of BtuB and TolC, low colE1-T-TolC affinity(*55*), and residually unblocked pore, even in the bound state. Because of their limited potency, our colE1-T and colE1-TR fragments are not likely to be practical in direct applications of antibiotic potentiation. However, our proof-of-concept findings offer a potential roadmap for further development. A more potent TolC binder would not need the R domain anchor for affinity. Such a binder could be designed using the atomic details of the colE1-T-TolC structure as a basis. Moreover, this scaffold can be used for efflux pump inhibitors for at least five other bacterial organisms. Each of these organisms has a structurally characterized outer membrane efflux pump that is homologous and structurally similar to TolC(*56*).

More broadly, there are at least nine known outer membrane protein receptors for bacteriocins. These outer membrane proteins have a variety of functions including adhesion, iron transport, and general import(*12, 57*). Using this same strategy with fragments derived from other bacteriocins may additionally allow for controlled inhibition of other bacterial functions.

## Materials and Methods

### *E. coli* strains

*E. coli* strains BW25113 and JW5503-1 were purchased from the Coli Genetic Stock Center (CGSC). JW5503-1 is a tolC732(del)::kan from the parent strain BW25113. BL21(DE3) were used for expression of the colicin constructs and TolC. BL21(DE3) has a premature stop codon at residue 58 of the *btuB* gene and therefore we used it as a Δ*btuB* strain for microscopy.

### Expression and Purification

The gene for colE1-TR was synthesized as a gBlock (Integrated DNA Technologies) and cloned into pET303. Inverse PCR was used to delete the R domain and produce colicin E1-T. The gene for colicin E1-TR-GFP was produced by inserting GFP at the C terminus of colE1-TR.

Plasmids were transformed into *E. coli* BL21(DE3) cells and plated on LB + agar + 100 μg/mL carbenicillin. Single colonies were inoculated into 50 mL LB broth with 100 μg/mL carbenicillin and grown overnight at 37 °C with shaking at 250 r.p.m. Proteins were expressed by inoculating 1L of TB supplemented with 0.4% glycerol, 10 mM MgCl_2_ and 100 μg/mL carbenicillin with 20 mL of the overnight culture. The culture was grown at 37 °C to an OD600 of 2.0 and induced with 1 mM IPTG. Expression cultures were then grown at 15 °C for 24 hours and harvested at 4,000 g for 30 minutes at 4 °C. Cell pellets were resuspended at 3 mL/g of cell pellet in lysis buffer (TBS, 5 mM MgCl_2_, 10 mM imidazole, 1mM PMSF, 10 μg/mL DNase, 0.25 mg/mL lysozyme) and lysed via sonication (2 minutes, 2s on, 8s off, 40% amplitude, QSonica Q500 with 12.7 mm probe) in an ice bath. Lysates were centrifuged at 4,000 g for 10 minutes to remove un-lysed cells and debris. The supernatant was centrifuged again at 50,400 g in a Beckman Coulter J2-21 for 1 hour at 4 °C. Clarified lysates were applied to a 5 mL HisTrap FF column and purified using an ÄKTA FPLC system with a 20 column volume wash step with binding buffer (TBS, 25 mM imidazole) and eluted using a linear gradient from 0-50% elution buffer (TBS, 500 mM imidazole) in 10 column volumes. Concentrated proteins were loaded onto a HiLoad Superdex 16/60 200 pg gel filtration column and eluted into phosphate buffered saline (PBS) pH 7.4.

TolC expression and purification was conducted in the same manner for preparation for cryoEM and for SEC. The gene for full-length TolC (a generous gift from R. Misra) was cloned into pTrc99a with the signal sequence deleted for expression into inclusion bodies. Plasmids were transformed into BL21(DE3) and plated on LB + agar + 100 μg/mL carbenicillin. A single colony was picked and grown in LB at 37 °C with shaking at 250 r.p.m. overnight. In the morning, 1L of LB was inoculated with 20 mL of the overnight culture and grown at 37 °C with shaking at 250 r.p.m. until the culture reached an OD600 of 0.6, at which point protein expression was induced with 1mM IPTG for an additional 4 hours. Cells were then harvested at 4,000g for 30 minutes at 4 °C. Cell pellets were resuspended in mL of lysis buffer (TBS, 5 mM MgCl_2_, 0.2 mg/mL lysozyme, 5 μg/mL DNase, 1mM PMSF) at 3 mL/1g of cell pellet and lysed via sonication (4 minutes, 2s on, 8s off, 40% amplitude, QSonica Q500 with 12.7 mm probe) in an ice bath. Cell lysates were centrifuged at 4,000 g for 30 minutes at 4 °C to harvest inclusion bodies. Inclusion body pellets were resuspended in inclusion body wash buffer (20 mM Tris pH 8.0, 0.5 mM EDTA, 1% Triton X-100) and centrifuged at 4,000g for 30 minutes at 4 °C. The inclusion body wash was repeated two more times. A final wash was done in 20 mM Tris pH 8.0 and inclusion bodies were stored at -20 °C. Inclusion bodies were solubilized in 20 mM Tris pH 8.0, 8M urea at 500 μM. N-octylpolyoxyethylene was added to 5 mL of solubilized TolC to a final concentration of 10% detergent and pipetted into a Slide-A-Lyzer G2 dialysis cassette with a 10,000 molecular weight cut off. Refolding was initiated by dialysis in 5L of 20 mM Tris pH 8.0, 100 mM NaCl at 4 °C with stirring overnight. TolC was centrifuged at 15,200 g for 10 minutes at 4 °C to remove aggregates. The supernatant was filtered through a 0.22 μm filter, concentrated to 2 mL, applied onto a HiLoad 16/60 Superdex 200 pg column on an ÄKTA Pure FPLC system, and eluted with 1.5 column volumes in 20 mM Tris pH 8.0, 100 mM NaCl, 0.05% n-dodecyl-β-D-maltoside. TolC containing fractions were pooled and concentrated to 300 μM in Amicon centrifugal filters with molecular weight cutoff of 30kDa.

Membrane scaffold protein MSP1D1. MSP1D1 in pET28a was purchased from Addgene and was expressed and purified as previously described(*58*).

### Peptide Synthesis

ColE1-T100-143 was synthesized using standard Fmoc chemistry with a CEM liberty blue microwave peptide synthesizer. The peptides were cleaved using a solution of 92.5:2.5:2.5:2.5 TFA:TIPS:H2O:DoDt and the crude peptides where purified using HPLC. Analytical HPLC traces were acquired using an Agilent 1100 quaternary pump and a Hamilton PRP-1 (polystyrene-divinylbenzene) reverse phase analytical column (7 μm particle size, 4 mm x 25 cm) with UV detection at 215 nm. The eluents were heated to 45 °C to reduce separation of rotational isomers, and elution was achieved with gradients of water/ acetonitrile (90:10 to 0:100 containing 0.1% TFA) over 20 min. Low-resolution mass spectra (LRMS) were obtained using a Waters Micromass ZQ 4000 instrument with ESI+ ionization

### Extracellular Protease Digestion

Protein localization after exogenous protein addition to whole cells was determined as previously described (*39*) with one modification. Samples for intact cells were lysed prior to loading on SDS-PAGE by adding 0.2 mg/mL lysozyme and incubating for 15 minutes followed by five freeze thaw cycles in liquid nitrogen.

### Single-Molecule Microscopy

Cysteine mutations were introduced at the C-terminus before the histidine tag for fluorophore conjugation. These constructs were purified as described in the expression and purification section with the addition of 1 mM TCEP in all buffers. All subsequent steps were performed with limited exposure to light and in amber tubes. Cyanine3 (Cy3) maleimide (Lumiprobe) was reconstituted in DMSO. Fluorophore labeling was achieved by mixing a 20-fold molar excess of Cy3 maleimide to protein and incubating overnight at 4 °C. Free dye was removed by gel filtration on a Sephadex NAP-10 G-25 column. Simultaneously to the dye removal, the sample was buffer exchanged into storage buffer (PBS pH 7.4, 1 mM DTT, 1 mM EDTA). The degree of labeling was determined spectrophotometrically from the concentrations of the dye and protein solutions using their respective extinction coefficients, ε, as described by their manufacturers or for the proteins as estimated by Expasy ProtParam (Cy3 ε548nm = 162,000 L mol-1 cm-1; colE1-T-E192C ε280nm = 9,970 L mol-1 cm-1; colE1-TR-E366C ε280nm = 14,440 L mol-1 cm-1). Labeling efficiencies were ∼75% and ∼85% for colE1-T-E192C and colE1-TR-E366C, respectively. Protein concentrations were adjusted according to the percentage of labeled protein.

Cultures of *E. coli* (WT, ΔtolC, or BL21(DE3)) were grown in LB medium at 37 °C with shaking (180 r.p.m.) overnight, then transferred to MOPS minimal medium (Teknova) with 0.2% glycerol and 1.32 mM K_2_HPO_4_, and grown at 37 °C for 13 h. The sample was transferred to MOPS medium and grown to turbidity at 37 °C overnight. A 1-mL aliquot of culture was centrifuged for 2 min at 4,850 g to pellet the cells. The pellet was washed in 1 mL MOPS and centrifuged a second time. The supernatant was then removed, and the cell pellet was resuspended in 500 μL MOPS. A 1.0 μL droplet of concentrated cells was placed onto a glass slide. Then, a 1.0 μL droplet of 1 µg/mL colicin E1 protein construct stock was added to the cells. The droplet was covered by an agarose pad (1% agarose in MOPS media) and a second coverslip.

Samples were imaged at room temperature using wide-field epifluorescence microscopy with sensitivity to detect single dye molecules as described previously (*40*). Briefly, fluorescence was excited by a 561-nm laser (Coherent Sapphire 560-50) for Cy3 or a 488-nm laser (Coherent Sapphire 488-50) for GFP. The lasers were operated at low power densities (1 – 2 W/cm2), and fluorescence was imaged with an Olympus IX71 inverted microscope with a 100x, 1.40-NA phase-contrast oil-immersion objective and appropriate excitation, emission, and dichroic filters. A Photometrics Evolve electron multiplying charge-coupled device (EMCCD) camera with > 90% quantum efficiency captured the images at a rate of 20 frames per second. Each detector pixel corresponds to a 49 nm × 49 nm area of the sample. Phase-contrast images of cells were segmented to attain *E. coli* cell outlines using a custom MATLAB script (The MathWorks, Natick, MA).

### Co-elution

The interaction of TolC and colicin E1-T or -TR were determined by co-elution on an SEC column. Purified TolC and colicin E1-T or -TR were mixed at a 1:2 molar ratio to ensure an excess of colicin to saturate TolC binding sites and incubated at room temperature for 1 hour before loading onto a Superdex 200 Increase 10/300 GL column (GE Healthcare). The protein was eluted with 1.5 column volumes into 20 mM Tris pH 8.0, 40 mM NaCl, 0.05% n-dodecyl-β-D-maltoside for colE1-T. For colE1-TR the NaCl concentration was increased to 200 mM to prevent precipitation. Elution fractions were collected every 0.5 mL. Peak fractions were concentrated to 20 μL and analyzed by 4-20% SDS-PAGE.

### Real-Time Efflux

Real-time efflux activity in the presence of colE1-TR was determined as previously described with some modifications(*48, 49*). Cells were resuspended to OD600 1.5 in cold PBS with and without 10-100 μM colicin proteins and incubated for 15 minutes on ice. To turn off efflux, 100μM carbonyl cyanide m-chlorophenyl hydrazone (CCCP) was added. After an additional 15 minutes the efflux dye NNN was added to the cells to 10 μM. The cells were incubated at 25 °C with shaking at 140 r.p.m. for 2 hours. Cells were harvested at 3,500 g for 5 minutes and washed once in 20 mM potassium phosphate buffer pH 7 with 1mM MgCl_2_. Cell concentrations were adjusted to OD600 1.0 and placed on ice. Then, 2 mL of the cell suspension was loaded into a quartz cuvette (Agilent Technologies). Fluorescence was measured with an Agilent Cary Eclipse fluorescence spectrophotometer with slit widths at 5 and 10 nm for excitation wavelength of 370 nm and emission wavelength of 415 nm. Fluorescence measurements were taken every 1 second. After 100 seconds, 50 mM glucose was added to re-energize the cells and initiate efflux, and fluorescence data were collected for an additional 600 seconds. Figure 5A-C, reflects time after glucose addition.

### Minimum Inhibitory Concentrations (MICs)

MICs were determined using the broth dilution method(*59*) with some modifications in 96 well plate format using LB media in 100 μL well volumes. Cultures were grown at 37 °C with shaking at 250 r.p.m. and OD600 was read on a Biotek plate reader after 20 hours. MICs are defined by the lowest concentration that prevents visible growth. We chose an OD600 of >0.1 as the cutoff for growth. We report MICs as the mean of 3 or 6 biological replicates with each replicate plotted (Figure 3D, Table S2). Due to the 2-fold discretized nature of concentration ranges used to determine MICs we do not report statistical significance values as is typical of MIC reporting.

### Reconstitution of TolC into Amphipol

Amphipol A8-35 (Anatrace) was solubilized in water at 33 mgs/mL. 1 mL of TolC at 0.5 mg/mL was mixed with 0.75 μL of A8-35 at 33 mg/mL for a 5- fold weight excess and incubated at room temperature for 30 minutes. Bio-Beads SM-2 resin that was washed in methanol and equilibrated with 20 mM Tris, 40 mM NaCl was added to the reaction mixture at 0.5 g/mL to initiate detergent exchange for A8-35 and incubated with rotation at 4 °C overnight. The mixture was transferred to a tube with fresh Bio-Beads and incubated at 4 °C for an additional 4 hours. The reaction mixture was loaded onto a HiLoad 16/60 Superdex 200 pg column on an ÄKTA Pure FPLC system and eluted with 1.5 column volumes in 20 mM Tris pH 8.0, 40 mM NaCl to remove free A8-35 and detergent. For colicin-bound TolC in A8-35, colicin E1-T was added to the reaction mixture at a >2 molar excess prior to gel filtration. TolC or colicin-bound TolC in A8-35 was concentrated to 2-4 mg/mL for cryoEM.

### Reconstitution of TolC into lipid nanodiscs

POPC (Avanti Polar Lipids) in chloroform was dried under a stream of nitrogen and freeze dried to remove residual chloroform. Lipids were reconstituted to 50 mM in cholate buffer (20 mM Tris pH 8.0, 100 mM NaCl, 0.5 mM EDTA, 100 mM cholate) in an amber glass vial. The vial was submerged under a stream of warm water until the solution became clear. Lipids, membrane scaffold protein, and TolC were mixed in a 36:1:0.4 ratio as previously described(*60*). Final concentrations were 4.5 mM POPC, 125 μM MSP1D1, 50 μM TolC in a 2 mL reaction with cholate brought up to 20 mM and dodecyl-maltoside up to 0.1%. The reaction mixture was incubated on ice for 1 hour. Bio-Beads SM-2 resin that was washed in methanol and equilibrated with 20 mM Tris, 40 mM NaCl were added to the reaction mixture at 0.5 g/mL to initiate nanodisc formation and incubated with rotation at 4°C overnight. The mixture was transferred to a tube with fresh Bio-Beads and incubated at 4 °C for an additional 4 hours. The reaction mixture was loaded onto a HiLoad 16/60 Superdex 200 pg column on an ÄKTA Pure FPLC system and eluted with 1.5 column volumes in 20 mM Tris pH 8.0, 40 mM NaCl to separate TolC inserted into nanodiscs from empty nanodiscs. For colicin-bound TolC in nanodiscs, colicin E1-T was added to the reaction mixture at a >2 molar excess prior to gel filtration. TolC or colicin-bound TolC in nanodiscs was concentrated to 2-4 mg/mL for cryoEM.

### Cryoelectron microscopy

3 μL of protein solution (TolC/colE1-T in amphipol or TolC/colE1-T in nanodiscs) was diluted to approximately 1.05 mg/mL concentration, applied to a glow-discharged TEM grid, and plunge-frozen in ethane using a Vitrobot Mark IV (FEI Company) with grade 595 filter paper (Ted Pella). Glow discharge was performed in ambient atmosphere at 0.39 mBar pressure. Imaging was performed using a Talos Arctica (FEI Company) operated at 200 kV with energy-filter and K2 Summit (Gatan, Inc.) for detection. To collect multiple images per hole while maintaining parallel illumination conditions, a nonstandard 20 μm condenser aperture was used to image TolC-colE1-T in nanodiscs. At nominal magnification of 205,000×, images were acquired in counting mode with a calibrated pixel size of 0.6578 Å. Fresnel fringes attributable to the beam interaction with the aperture were often seen in images. Some investigators have moved the microscope stage and altered the nominal objective lens true focus point to generate a fringe-free condition(*61*). In this study, imaging at 205,000× with a 20 μm aperture yielded better results than imaging at 130,000× with a 50 μm aperture; at 130,000× with a 20 μm aperture the fringes were extremely severe due to the larger field of view, so a full dataset was not collected with those conditions. TolC in nanodiscs (without colE1-T) was imaged at 130,000× with a 50μm condenser aperture (Table S1).

Micrographs were collected with SerialEM(*62*) using in-house modifications of the scripts of Chen Xu (sphinx-emdocs.readthedocs.io). Briefly, multishot imaging was configured with 4 images per hole for each of 16 holes; intermediate-magnification montages of grid squares were acquired; points were selected manually for collection of 64 images per point; images were acquired using coma-compensated image shift as gain- and dark-corrected LZW-compressed TIFs. Side, top, and oblique views were seen in areas of thin ice. During screening, ice thickness was estimated at 17-30nm by the method of *I*_0_/*I*_ZLP_(*63*).

The collection of micrographs of TolC without colicin at 130,000× magnification has been previously described(*64*).

### 3D reconstruction and modeling

Final reconstructions were obtained using cryoSPARC 2(*65*). 1,018 micrographs were collected of amphipol-embedded TolC/colE1-T. Micrographs were motion-corrected using RELION 3(*66*). CTF parameters were determined by means of *ctffind*(*67*). 115,362 particles were selected with crYOLO(*68*). 2D classification revealed that many particles had aberrant morphology and only 24,624 (21%) were selected for 3D reconstruction. *Ab initio* reconstruction in cryoSPARC 2(*65*) was effective at recovering a map whose shape was similar to that of previously described TolC trimers. However, the data could only be refined to a nominal global resolution of 6.0 Å, and lumenal density was insufficiently resolved. 4,492 micrographs were then collected of nanodisc-embedded TolC/colE1-T and processed similarly. Of the 339,779 particles detected by cryoSPARC Live, 179,834 (53%) were in good classes. Although there were slightly fewer particles per micrograph in the nanodisc dataset, more particles per micrograph were usable. Beginning with the *ab initio* model and mask derived from the amphipol data, this particle set was refined by cryoSPARC 2 non-uniform refinement with or without imposed C*_3_* symmetry. The maps refined to nominal global resolutions of 2.81 Å and 3.09 Å for the symmetrized and asymmetric maps, respectively. There was local variation in resolution within the map, with consistent, high resolution in the middle of TolC and lower resolution at the lids and for colE1-T. Local resolution was computed in cryoSPARC by the locally windowed FSC method(*69*) and rendered with UCSF Chimera. To reduce the voxel-based values to averages for four regions of the complex, the local resolution map was masked to include only voxels within 3 Å of a modeled atom and then the median value was calculated for those voxels closest to colE1, closest to TolC residues 168-172 and 386-390, closest to TolC residues 285-301 and 76-82, or closest to other TolC residues. Furthermore, it is notable that while the nanodisc appears as a double-belt in the symmetrized map, in the asymmetric map the nanodisc protein mostly appears on the side of TolC that is bound to colE1-T. One possible explanation is that, despite masking, nanodisc asymmetry is a confounder of the asymmetric refinement and is one source of heterogeneity in the data. Another possibility is that the C-terminus of colE1-T forms an interaction with the nanodisc, causing preferential alignment of the nanodisc with respect to the TolC/colE1-T complex.

196,158 particles of TolC without colicin were obtained as previously described(*64*). Homogeneous refinement yielded a structure at 2.89 Å; local motion correction and global CTF refinement yielded a final map at 2.84 Å.

Modeling was initiated by rigidly docking a crystal structure of TolC in complex with hexamminecobalt (1TQQ)(*31*) into the symmetrized map density. Automated, semi-automated, and manual real-space refinement was performed using phenix(*70*), ISOLDE(*71*), and coot(*72*). For TolC with colE1-T, additional refinement was performed in AMBER using the cryoEM density map as a restraint.

Although additional residues are present at the TolC C-terminus, these were not modeled because the density becomes unsharp after residue 428. Blurred density in the map suggests that the C-terminus follows helix 3 towards the periplasmic end of the molecule. ColE1-T was modeled *ab initio*. The 3-fold symmetrized map contains density at ∼1/3^rd^ occupancy for colE1-T and this density contains some high-resolution information not present in the asymmetric map, except near the TolC lid regions where symmetrization overlays colE1-T density with TolC density at a threefold-related position. First, polyalanine helices were placed in the helical density in coot.

Cross-correlation coefficients for both helices are higher with the N-terminus oriented towards the periplasm, and the Christmas tree appearance was observed indicating that this is the correct chain orientation. An estimate of the registration was made by visual inspection of potential anchor residues. Finally, the hinge region was filled in using phenix and coot. This completed chain was refined against the asymmetric map in ISOLDE. Iteration between phenix, coot, and ISOLDE was continued until acceptable fit to density was achieved. In the case of TolC with colE1-T, the map was further improved by combining map information with molecular dynamics force fields(*34*). Briefly, starting with the phenix/coot/ISOLDE-refined model, we performed restrained simulated annealing in AMBER, heating from 0K to 300K for 0.2 nsec, holding constant temperature for 1 nsec, and then cooling to 0K over 0.2 nsec. The cryoEM density map is utilized as a restraint potential in the annealing so that both map information and AMBER force field information are simultaneously utilized to obtain an optimum model consistent with the data(*34*). The protein force field used the ff14SB force field(*73*) and a generalized Born implicit solvent model with *igb=8*(*74*), and a nonbonded cutoff of 20 Å. The relative weight of real-space map-based restraints and the force field was fixed using *fcons=0.02*. For colE1-T, information from the symmetrized map was integrated into the modeling procedure during manual remodeling in coot, but map-based refinement in phenix, ISOLDE, and AMBER were against the asymmetric map. TolC without colE1-T was modeled similarly but using the TolC-colE1-T structure as a starting point instead of 1TQQ, and without final AMBER refinement.

Molecular representations were generated with Chimera, ChimeraX(*75*) or PyMOL (Schrödinger, LLC).

## Acknowledgments

We gratefully acknowledge Daniel Montezano, Pinakin Sukthankar, Rik Dhar, Dwight Deay, Scott Lovell, Matthias Wolf, Alexander Little, Heather Shinogle, and Sarah Noga for discussions and feedback, Mark Richter for the use of his fluorometer, Rajeev Misra for the pTrc vector containing the TolC gene, Vasileios Petrou for guidance on nanodiscs, Chamani Perera for peptide synthesis, Karen Marom for editorial guidance. We thank the Office of Advanced Research computing (OARC) at Rutgers for high-performance computing resources.

## Funding

NIH award R21-GM128022 to JSB

NIGMS awards DP2GM128201, P20GM113117, P20GM103638 and the Gordon and Betty Moore Inventor Fellowship to JSGS,

NIGMS awards P20 GM103418 and 2K12GM063651 to SJB.

## Author contributions

Conceptualization: SJB, JSGS Methodology: SJB, JTK, JSGS, JSB

Investigation: SJB, JJS, ALC, API, VKW, EF, DAC, JTK

Writing—original draft: SJB, JSGS Writing—review & editing: SJB, JSGS, JSB, JTK

## Competing interests

Authors declare that they have no competing interests.

## Data and materials availability

All data are available in the main text or the supplementary materials.” All strains used are commercially available. All plasmids available through Addgene. CryoEM maps and models have been deposited with accession codes EMD-21960, EMD-21959, PDB ID 6WXI, and PDB ID 6WXH. All other data is available in the main text or the supplementary materials.

## Supplementary Materials

**Figure S1.**
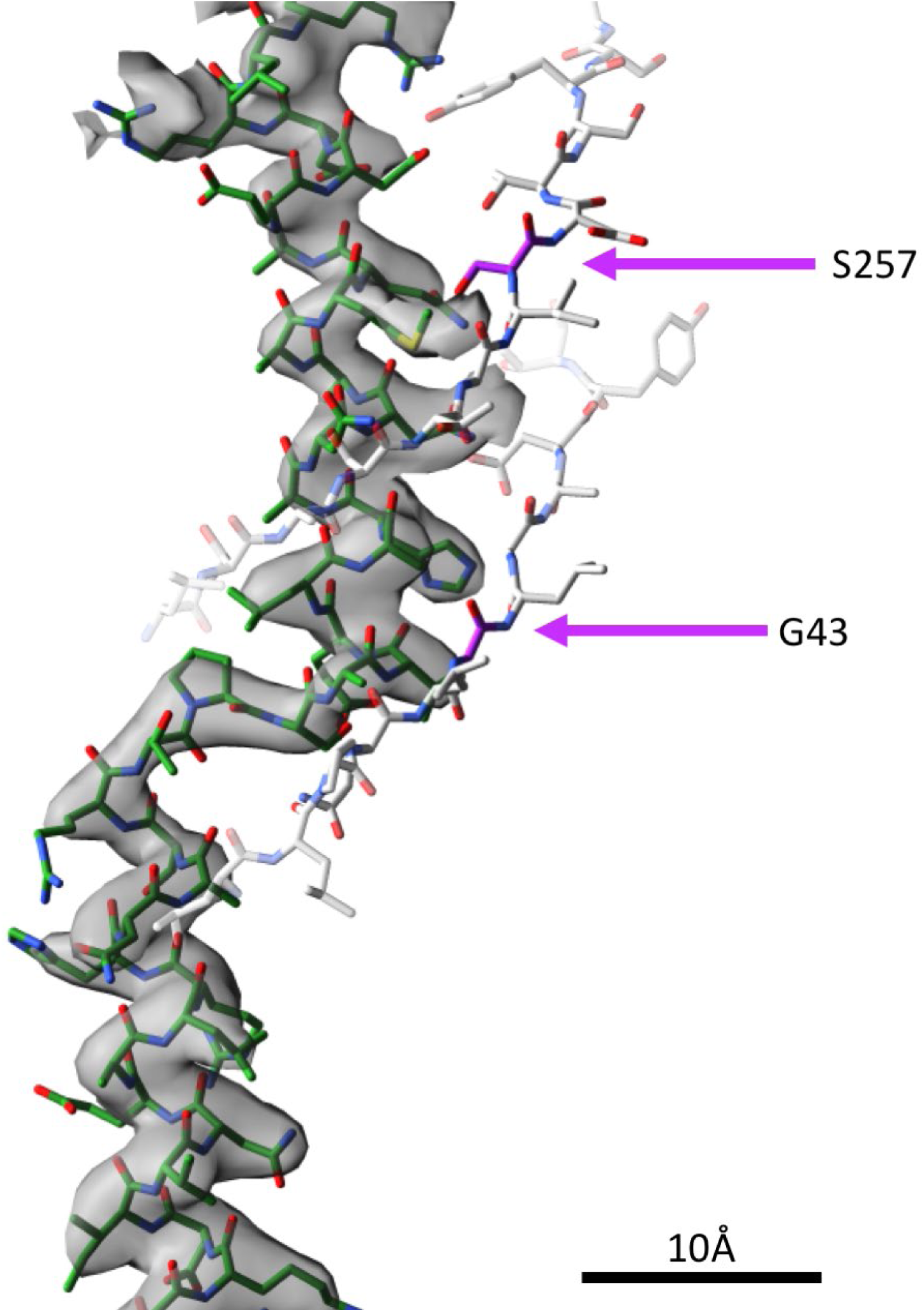
TolC box density. cropped from the fully asymmetric cryoEM map of the TolC/colE1-T complex. Mutations at positions G43 and S257 in the TolC (grey sticks) barrel make direct contact with colE1-T (green). These mutations were previously shown to abolish WT colE1 function(*37*). Mutations to bulkier side chains introduce a steric clash that prevents binding.

**Figure S2.**
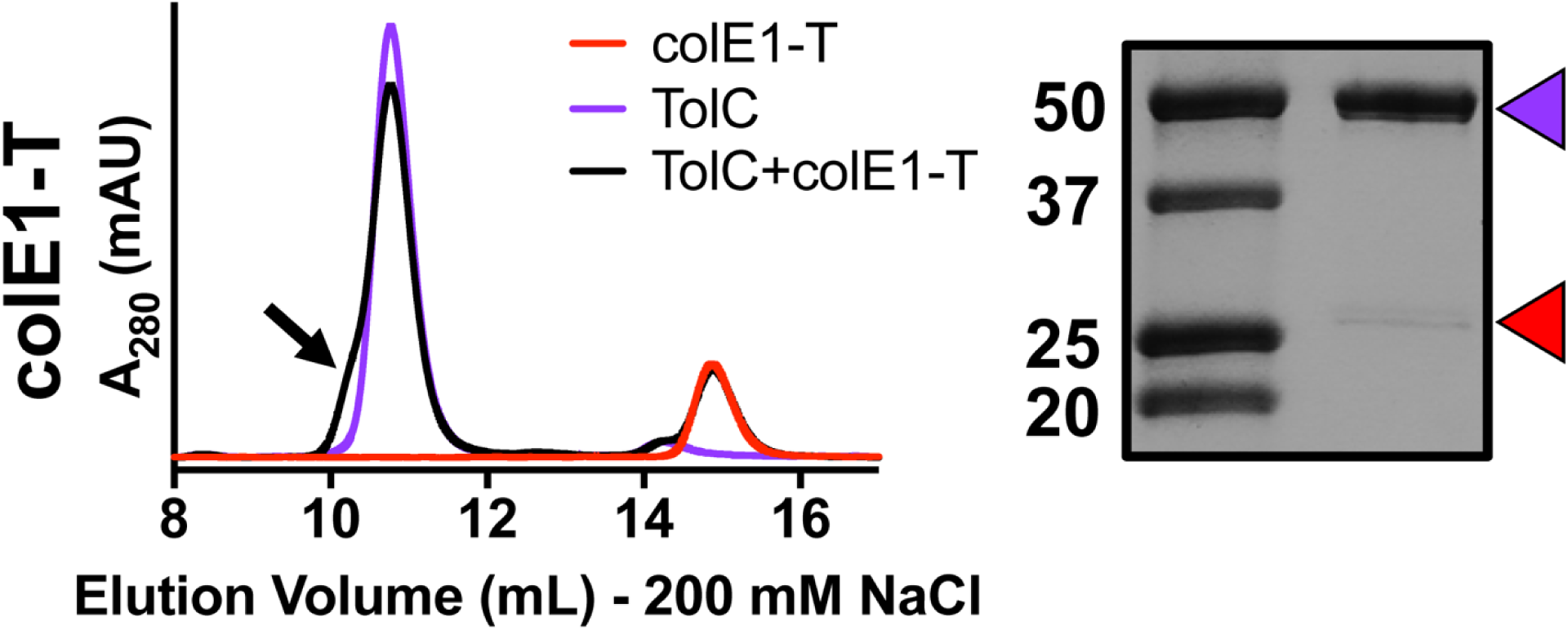
Co-elution of colE1-T with TolC at 200 mM NaCl (black). Under this higher salt concentration, when TolC (purple) and colE1-T (red) are mixed, there is a smaller peak shift than that seen with colE1-TR and the presence of a shoulder (black arrow). SDS-PAGE of the shoulder shows presence of both TolC (purple arrow) and colE1-T (red arrow). Although some binding was detected, this higher salt concentration prevents full binding as indicated by a much fainter band for colE1-T than seen at the lower salt concentration (Figure 4A).

**Figure S3.**
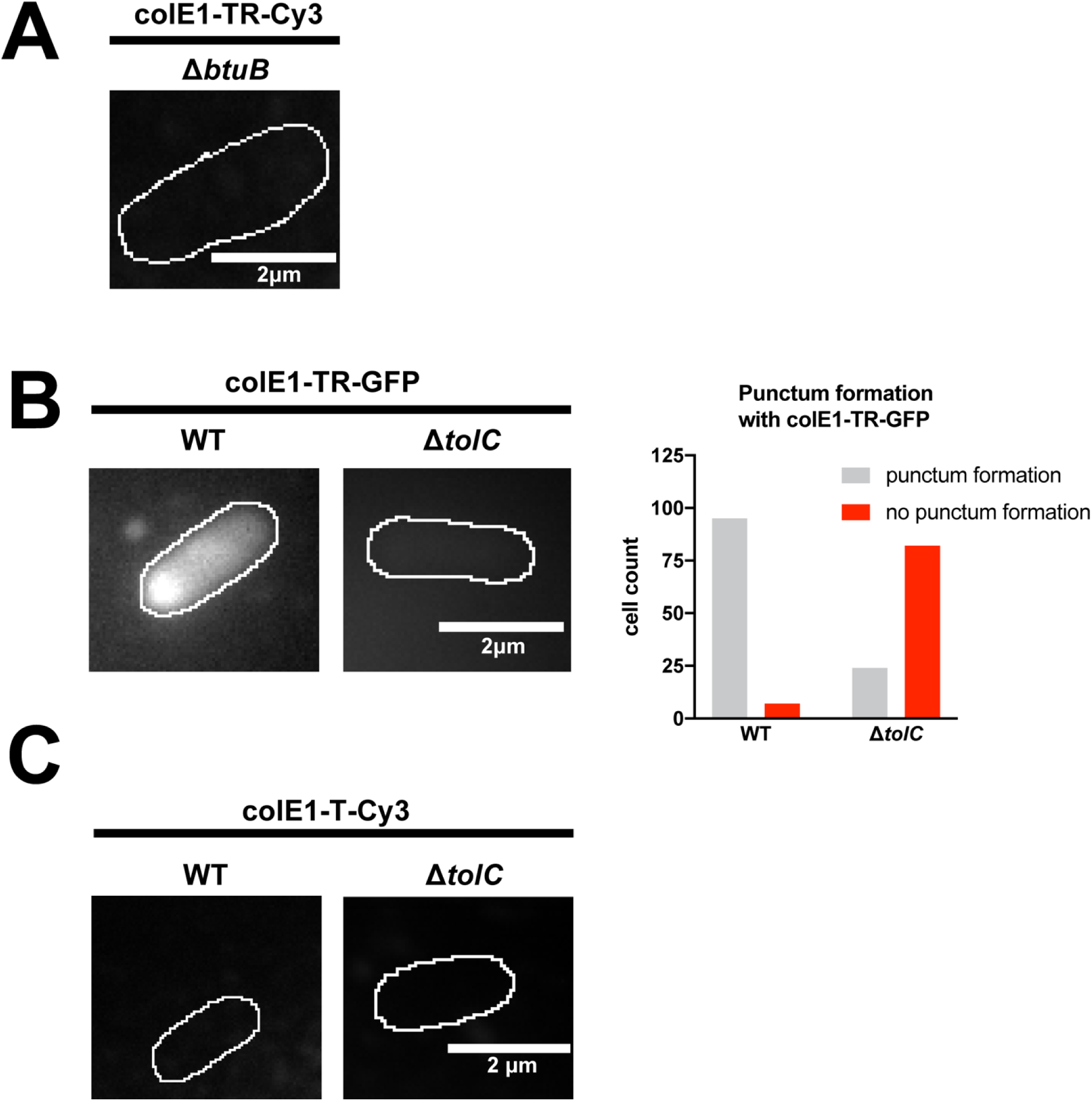
Single-molecule microscopy. Fluorescence images overlaid on outlines of living *E. coli* cells from phase-contrast microscopy for WT and Δ*tolC* for colE1-TR-GFP and counts of cells where colE1-TR-GFP punctum formation was observed. (A) 97% of cells showed no binding of Cy3-labeled colE1-TR to Δ*btuB* (B) ColE1-TR-GFP forms similar puncta as Cy3-labeled ColE1-TR (main text Figure 4C). (C) No binding of Cy3-labeled colE1-T to WT or Δ*tolC* cells was detected with Cy3-labeled colE1-T. Scale bars: 2 µm.

**Figure S4.**
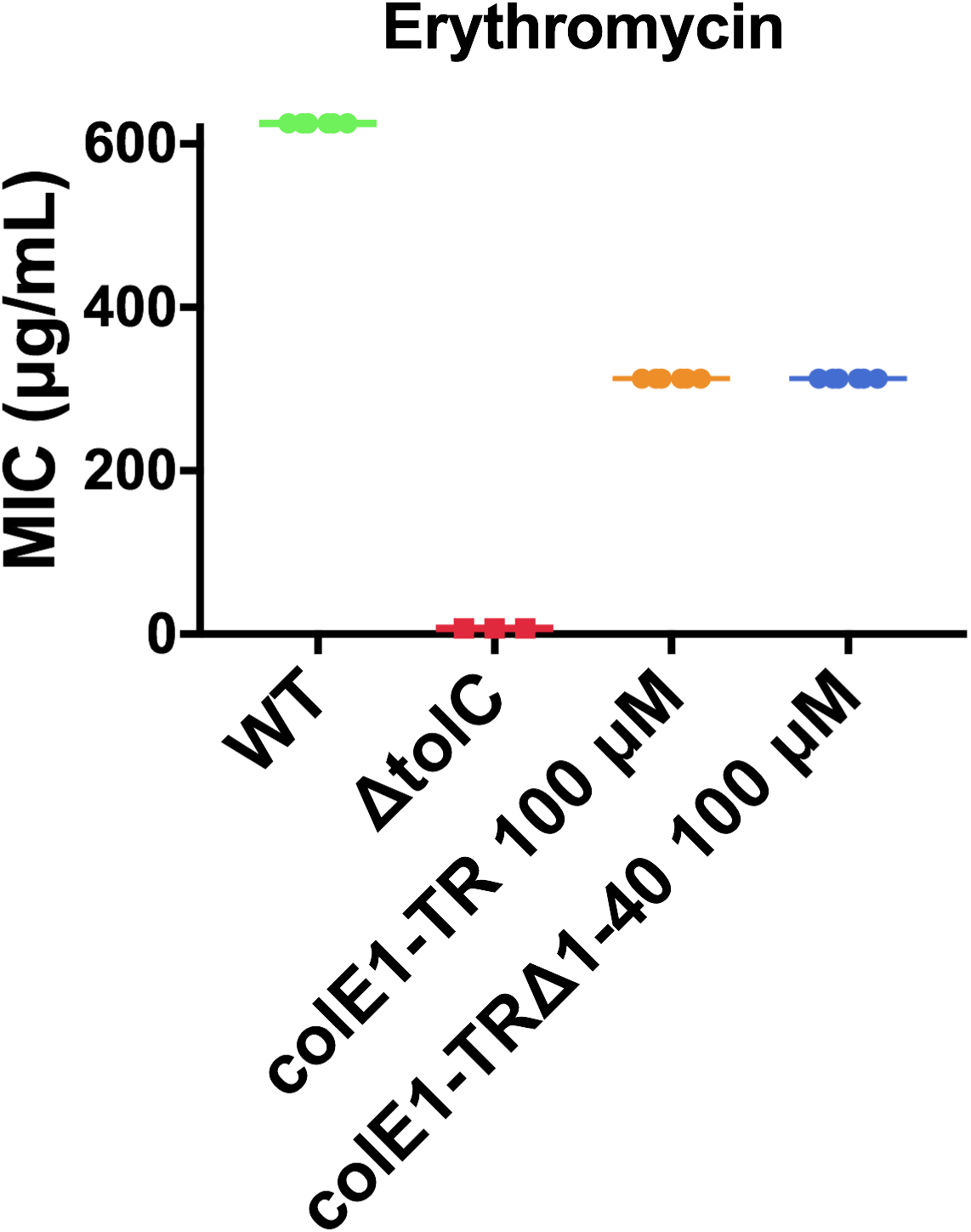
Minimum Inhibitory Concentration of Erythromycin with colE1-TR TolA box deletion construct. The MIC for erythromycin with 100 μM colE1-TR or colE1-TR_Δ1-40_ is identical indicating that TolA engagement by colE1-TR does not contribute to antibiotic susceptibility.

**Table S1.** Cryo-EM data collection, refinement, and validation statistics

|  | TolC<br>(EMDB-21960)<br>(PDB 6WXI) | TolC + colE1<br>(EMDB-21959)<br>(PDB 6WXH) |
| --- | --- | --- |
| Data collection and processing |  |  |
| Electron microscope | Talos Arctica | Talos Arctica |
| Magnification | 130,000× | 205,000× |
| Voltage (kV) | 200 | 200 |
| Electron exposure (e-/Å<sup>2</sup>) | 33.04 | 35.96 |
| Defocus range (μm) | 0.5-2.6 | 0.4-2.1 |
| Pixel size (Å) | 1.038 | 0.6578 |
| Symmetry imposed | C<sub>3</sub> | C<sub>1</sub> (C<sub>3</sub>) |
| Initial particle images (no.) | 2,092,678 | 339,779 |
| Final particle images (no.) | 778,220 | 179,834 |
| Map resolution (Å) | 2.84 | 3.09 (2.81) |
| FSC threshold | 0.143 | 0.143 |
| Refinement |  |  |
| Initial model used (PDB code) | 6WXH | 1TQQ |
| Model resolution (Å) | 3.11 | 3.37 |
| FSC threshold | 0.5 | 0.5 |
| R.m.s. deviations |  |  |
| Bond lengths (Å) | 0.0072 | 0.012 |
| Bond angles (°) | 1.11 | 1.76 |
| Validation |  |  |
| MolProbity score | 1.33 | 0.50 |
| Clashscore | 4.42 | 0.00 |
| Poor rotamers (%) | 1.40 | 0.18 |
| Ramachandran plot |  |  |
| Favored (%) | 98.12 | 98.31 |
| Allowed (%) | 1.88 | 1.32 |
| Disallowed (%) | 0 | 0.37 |

**Table S2.** Mean minimum inhibitory concentrations (MICs) of antimicrobials in the presence of colE1-TR.

|  | Erythromycin<br>(µg/mL) | Ciprofloxacin<br>(ng/mL) | Benzalkonium<br>Chloride (µg/mL) |
| --- | --- | --- | --- |
| WT | 625.00 | 56.25 | 50 |
| *ΔtolC* | 7.03 | 10.16 | 4.125 |
| WT + 10<br>µM<br>colE1-TR | 312.50 | 75.00 | 50 |
| WT +<br>100 µM<br>colE1-TR | 312.50 | 28.13 | 25 |

**Movie S1.**

A representative 8-second movie of colE1-TR bound to a live cell. ColE1-TR localizes on, and remains bound to, the extracellular surface of *E. coli*. Fluorescence movie of Cy3-labeled colE1-TR on living WT *E. coli* overlaid on outline of the *E. coli* cell from phase-contrast microscopy. Continuous imaging at 25 frames per second; the movie plays in real time at the same speed. This movie corresponds to the first 8 seconds of data used to attain the WT image in Figure 4C. Scale bar: 2 µm.

## Notes

### Competing Interest Statement

The authors have declared no competing interest.

